# Structure-activity relationship of an all-α-helical prenyltransferase reveals a mechanism for indole prenylation

**DOI:** 10.1101/2025.05.02.651859

**Authors:** Takumi Oshiro, Shuta Uehara, Yoshikazu Tanaka, Takuya Ito, Yoshio Kodera, Takashi Matsui

**Affiliations:** Department of Physics, School of Science, Kitasato University, Kanagawa 252-0373 Japan; Graduate School of Life Science, Tohoku University, Miyagi 980-8577 Japan; Faculty of Pharmacy, Osaka Ohtani University, Osaka 585-8540 Japan; Center for Disease Proteomics, School of Science, Kitasato University. Kanagawa 252-0373 Japan

**Author notes:** Correspondence: Dr. Takashi Matsui, Postal address: 1- 15-1 Kitasato, Minami-ku, Sagamihara, Kanagawa 252-0373, Japan.

## Abstract

Enzymes are involved in the biosynthesis of various secondary metabolites found in nature. The catalytic mechanism is regulated by the three-dimensional structure of the enzyme especially at the catalytic site, resulting in the natural products with complicated conformation derived from a regioselective, chemoselective, and stereoselective fashion of the enzyme reaction. Prenyltransferase (PT), which belongs to the prenylsynthase (PS) family, catalyzes the condensation of isoprene to an aromatic compound, consequently producing a terpenoid scaffold structure. Therefore, it plays an important role in which expands the chemical diversity of terpenoids. Although the three-dimensional structures of PS which categorized in the same family were resolved, the catalytic mechanism of the PT has been vailed. In this study, we determined the X-ray crystal structure of a novel prenyltransferase *Sx*PT1 derived from marine *Streptomyces*. Here we described that *Sx*PT1 analyzed the enzyme reactions and discussed its catalytic mechanism.

## Introduction

With more than 440,000 natural compounds identified to date, these compounds are ubiquitous across all domains of life and exhibit a wide range of biological activities^1,2^. Terpenoids, which represent the largest and most widely distributed family of natural products^3^, play important ecological roles in nature, such as maintaining membrane stability, facilitating photosynthesis, and providing chemical defense mechanisms^4^. Furthermore, they are widely used in human life such as a perfume, pharmaceuticals, and fuels^3,5,6^. This enormous variety derived from diverse terpene scaffolds is involved in the enzymes in the prenylsynthase family including synthases, cyclases and transferases. In primary metabolism, a prenylsynthase condensates an isopentenyl diphosphate (IPP, C_5_) with a dimethylallyl diphosphate (DMAPP, C_5_), consequently generating a geranyl diphosphate (GPP, C_10_). Some prenylsynthases accommodate longer chains of isoprenoids to condensate additional rounds of IPP to elongates isoprene units and generate the longer isoprenoids such as farnesyl diphosphate (FPP; C_15_) and geranylgeranyl diphosphate (GGPP; C_20_)^7–9^. Some isoprenoids produced by the prenylsynthase are cyclized by prenylcyclase to form compounds used in perfumes, e.g., γ-terpinene^10^. The three-dimensional structures and catalytic mechanisms of prenylsynthase and prenylcyclase are well known^11–13^. They entirely consist of α-helices and share similar catalytic mechanisms^8,11^. Additionally, they contain highly conserved metal ion coordinating domains (DDXXD), referred to as the aspartate-rich motif (ARM)^8,14^.

Other isoprenoids are often serve as starter substrates to produce various indole alkaloids. PaxC, an ABBA type indole prenyltransferase, condensates GGPP to an indole-3-glycerol phosphate to give a key intermediate to produce paxillin. In addition, ABBA type enzymes exhibit a promiscuous substrate recognition, generating a wide variety of products, and share a similar antiparallel αββα barrel conformation^15^. Structure-based engineering of the ABBA type enzyme has successfully increased substrate acceptance^16^ and facilitated the production of a diverse range of novel compounds by providing different substrates^17^. UbiA, another type of membrane-embedded prenyltransferase, accommodates a broad range of substrates, ranging from DMAPP (C_5_)^18^ to dodecaprenyl phosphate (C_60_)^19^, as a prenyl donor substrates and is involved in the biosynthesis of ubiquinone^20,21^. UbiA type enzyme has two characteristics conserved motifs, NDXXDXXXD and DXXXD^22^, similar to those found in all-α-helical prenylsynthase. Three-dimensional structure of UbiA has revealed how the prenyltransferase reaction is carried out^18^. While, the synthase forms a soluble homodimer, UbiA is a membrane protein that presents as a monomer on the membrane.

According to the proposed biosynthetic mechanism of indolosesquiterpenoid, xiamycin A^23,24^, the other type of prenyltransferase would be proposed. This enzyme shows higher sequence similarity to the prenylsynthases and cyclases rather than other type of transferases, e.g., ABBA-type transferase, and shares an aspartate rich motif (ARM), suggesting it is a member of the all-α-helical prenylsynthase superfamily. Recently, *in silico* approach supplemented by AlphaFold2-informed protein model^25^ predicted that all α-helical prenyltransferase prenylates polypeptide through a dissociative electrophilic substitution mechanism^26–29^. Previous research has shown that all-α-helical type prenyltransferase possesses a first ARM (FARM), while the region corresponding to the second ARM (SARM) is less conserved and thus is named pseudo-SARM^30^. Furthermore, previous mutational studies have suggested that FARM and pseudo-SARM are important for enzyme reactions^25,30,31^. However, three-dimensional structure of the all-α-helical type prenyltransferase had remained elusive, and the mechanism of this type of prenylation had been still unclear.

Here, we focused on an all-α-helical prenyltransferase named *Sx*PT1 derived from *Streptomyces xiamenensis*. *Sx*IPT1 was proposed to function as a prenyltransferase due to the presence of its conserved FARM and pseudo-SARM, and high sequence homology to the xiamycin A biosynthesized enzyme XiaM ^23^. In this paper, we report the X-ray crystal structure of *Sx*PT1, and the prenylation mechanism involved in the biosynthesis of all-α- helical type prenyltransferase was discussed.

## Materials and methods

### Protein expression and purification of SxPT1and its mutants

The *SxPT1*gene was cloned into a pET–26b vector (Merck, Darmstadt, Germany). *Sx*PT1 fused with C-terminal His_6_ tag was overexpressed in *E. coli* strain Rosetta2 (DE3) (Merck) under the control of a T7 promoter. Cells harboring the plasmid were grown in an LB medium containing 20 mg/L kanamycin and 25 mg/L chloramphenicol at 37°C until the O.D_600_ reached 0.6. 0.125 mM isopropyl-β-D-1-thiogalactopyranoside (IPTG) was then added to induce *Sx*PT1, and growth was continued for 15 h at 20°C. The cells were harvested by centrifugation at 8,000×*g* for 15 min at 4°C and resuspended in 40 mL of lysis buffer (50 mM Tris-HCl pH 8.0, 300 mM NaCl, 10% (v/v) glycerol). The cell lysate was centrifuged at 8,000×*g* for 15 min at 4°C. The supernatant was pathed through a 5-μm filter device (Merck) and loaded onto a 1 mL of HisTrap HP (Cytiva, Marlborough, MA, USA) equilibrated with lysis buffer. The recombinant protein was eluted with lysis buffer containing 300 mM imidazole. The protein solution was further purified to homogeneity by size-exclusion chromatography on a HiPrep 16/60 Sephacryl S-200 (Cytiva) equilibrated with lysis buffer. Appropriate fractions were analyzed for purity using SDS-PAGE. The fraction containing *Sx*PT1 was concentrated to 12 mg/mL, measuring by the UV absorption at *A*_280_, with an Amicon Ultra 15 centrifugation device (Cytiva).

Selenomethionine labeled *Sx*PT1 (SeMet-labeled *Sx*PT1) was overexpressed in *E. coli* strain Rosseta2 (DE3). Cells were grown at 37°C in M9 medium containing 1 g/L NH_4_Cl, 1 mM MgSO_4_, 0.1 mM CaCl_2,_ 1×BME vitamins solution (Merck), 0.4%(w/v) glucose, 20 mg/L kanamycin and 25 mg/L chloramphenicol until the O.D_600_ reached 0.6. 25 mg/L selenomethionine, and 100 mg/L each of L-Lys, L-Thr, L-Ile, L-Leu, L-Val, and L-Phe were added and grown for 30 min at 37°C, subsequently 0.125 mM IPTG was added to express SeMet-labeled *Sx*PT1, and growth was continued for 15 h at 20°C. Purification of SeMet-labeled *Sx*PT1 was performed as described above.

### Site-direct mutagenesis

The plasmids expressing *Sx*PT1 mutant enzymes were constructed with a KOD-Plus-mutagenesis kit (TOYOBO, Osaka, Japan), according to the manufacturer’s protocol using *Sx*PT1 expression plasmid as a template. Tyr215 was substituted by Phe (to give *Sx*PT1 Y215F), Tyr215 was substituted by Ala (to give *Sx*PT1 Y215A), Gln216 was substituted by Ala (to give *Sx*PT1 Q216A), and Asp219 was substituted by Ala (to give D219A). The mutant enzymes were expressed and purified with the same procedures as described for the wild type of *Sx*PT1. All primers used to generate *Sx*PT1mutants can be found in Supplementary Table S1.

### Enzyme assay of SxPT1

30 mM of *Sx*PT1 or mutant enzymes was co-incubated with 1 mM FPP, 1 mM Imidazole and 5 mM MgCl_2_ in 100 μL of 100 mM Tris-HCl pH 8.0, at 20°C for 12 h. The enzyme reaction products were extracted with an equal volume of ethyl acetate. The organic layer was analyzed by HPLC (Nano Space SI-2, Shiseido, Tokyo, Japan). The enzyme reaction products were eluted with a gradient of buffer A (0.01% formic acid) and buffer B (90% CH_3_CN,0.001% formic acid): 0-6 min, 10% buffer B; 6-28 min, 10-100% buffer B, on a CAPCELL PAK C18 BB-H (2.1 mm I.D. × 100 mm; OSAKA SODA, Osaka, Japan) at a flow rate of 0.2 mL/min.

The Products were analyzed by LC-MS analysis using a quadrupole Orbitrap benchtop mass spectrometer Q-Exactive (Thermo Fisher Scientific) equipped with HPLC (Nanospace SI-2; Shiseido). The enzyme reaction products were injected directly into CAPCEL PAK C_18_ BB-H at a flow rate of 3 μL/min. The Enzyme reaction products were separated with a gradient of solvents A and buffer B: 0-2 min, 30% buffer B; 2-8 min, 30-100% buffer B; 8-18 min, 100% buffer B. The analytical conditions were followed previously described^32^ with slight modifications. MS1 spectra were collected in the scan range of 200–700 *m*/*z* at 140,000 resolution to hit an automatic gain control (AGC) target of 1×10^6^.

### Crystal structure determinations of *Sx*PT1 and their mutant enzymes

All crystallization attempts were performed using the sitting–drop vapor diffusion technique. Diffraction quality crystals of *Sx*PT1, SeMet-labeled *Sx*PT1, Q216A and D219A mutant enzymes were appeared after a few days at 18°C. All crystallization drops were prepared by mixing 0.5 μL of either the purified protein solutions and an equal volume of reservoir solution and equilibrating the mixtures against 50 μL of reservoir solution. To obtain a substrate complex structure, *Sx*PT1 or mutant enzyme crystal was soaked in reservoir containing 1 mM FSPP, 5 mM indole, and 5 mM MgSO_4_ for 24 h at 20°C.

The diffraction quality crystals were transferred into the reservoir solution with 10%(v/v) ethylene glycol and then flash cooled in a liquid nitrogen. X-ray diffraction data sets were collected at the BL-1A beamline at the Photon Factory (Tsukuba, Japan) for the crystals of *Sx*PT1 WT, *Sx*PT1-FSPP and *Sx*PT1 Q216A-FSPP and BL-5A beamline at the Photon Factory for the crystal of *Sx*PT1 SeMet and BL-17A beamline at the Photon Factory for the crystal of *Sx*PT1 D219A, respectively. Wavelength of 0.97922 Å for single-wavelength anomalous diffraction phasing method, and 1.04500 Å was used for data collection of *Sx*PT1 WT, *Sx*PT1-FSPP and *Sx*PT1 Q216A-FSPP, and 0.98000 Å was used for data collection of *Sx*PT1 D219A structures, respectively.

Diffraction data for single-wavelength anomalous diffraction method^33^ were processed and scaled using the XDS^34^ program package. Se sites were determined with Autosol in PHENIX^35^. The sites were refined, and the initial phases were calculated with AutoBuild in PHENIX^36^. The initial phases of the *Sx*PT1 apo structure were determined by molecular replacement, using the structure of SeMet-labeled *Sx*PT1 as a search model. Molecular replacements were performed with the *Sx*PT1 wild-type as the search model using Molrep in CCP4 suite^37^ to solve the complex structure with FSPP and these mutant enzymes. The structure was modified manually with COOT^38^ and refined with Phenix.Refine^39^, with the twin operators. The detailed data collection and refinement statistics are summarized in Supplementary Table S2. The quality of the final models was assessed with Molprobity^40^. All crystallographic figures were prepared with PyMOL^41^ and ChimeraX^42^.

## Results

### Structure of *Sx*PT1

To understand the structural basis for the substrate recognition, the initial phase of *Sx*PT1 was obtained at the resolution of 1.8 Å using Se single-wavelength anomalous dispersion (Se-SAD). The crystal structure of apo *Sx*PT1 was solved at 2.1 Å resolution using the molecular replacement method calculated from SeMet-labeled *Sx*PT1 as a template model. The crystal structure of apo *Sx*PT1 consists of two homodimers formed by molecules A-B and molecules C-D in the asymmetric unit (Figure 1A). The overall structure of *Sx*PT1 is composed of 16 α-helices. The molecules B, C, and D exhibited the root mean square deviation (RMSD) values of 1.0, 0.2, and 0.9 Å for the molecule A, respectively. Molecule B also converged well to molecule D with the rmsd value of 0.8 Å, suggesting that the homodimer composed of molecules A and B had similar overall conformation with the other homodimer composed of molecules C-D.

**Figure 1.**
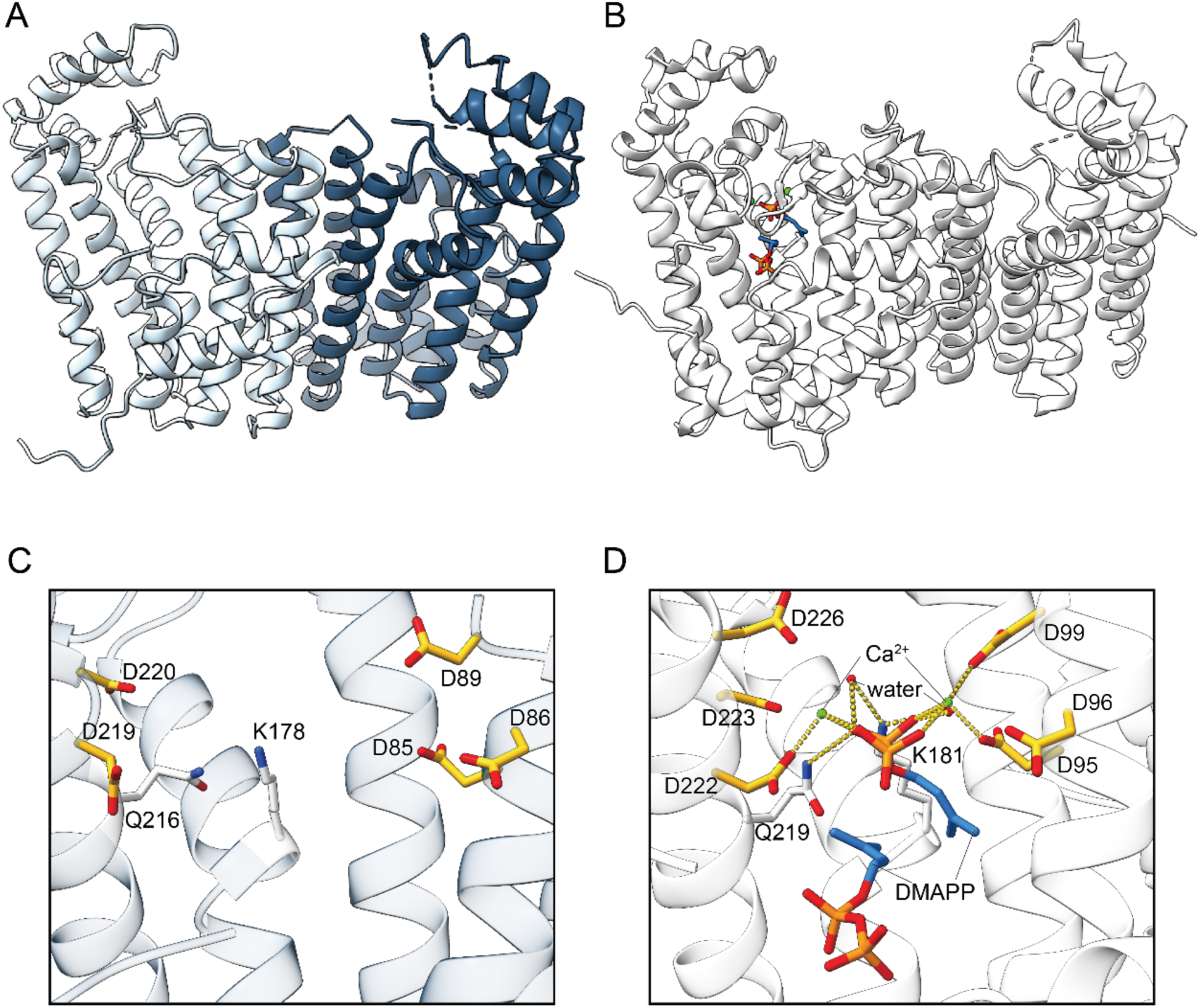
Crystal structure comparison of apo *Sx*PT1 with the prenylsynthase 3OYR. The overall structures of (A) *Sx*PT1 and (B) 3OYR (PDB ID: 3OYR) are illustrated as a ribbon diagram. The structure of *Sx*PT1 is colored by protomer. The comparison between the conserved ARMs in (C) *Sx*PT1 and (D) 3OYR. Sidechains of the aspartic acids in the motifs are shown as yellow sticks. The coordinations of divalent ions were indicated as yellow dotted lines.

The similar conformational enzymes explored using Dali^43^, several prenylsynthases, such as EcOPPs (PDB ID 3WJO, 26.6% identity with *Sx*PT1) and PATL_3739 (PDB ID 4JXY, 27.6% identity with *Sx*PT1), were assigned with the RMSD values of 2.4 Å (z-score 29.2) and 2.1 Å (z-score 29.4) for *Sx*PT1, respectively (Supplementary Figure S1). A prenylsynthase (PDB ID 3OYR, no unique name has been assigned, thus referred to as 3OYR in this paper, Figure 1B) also exhibited an RMSD value of 1.9 Å with a z-score of 28.9 for *Sx*PT1 coupled with highest z-socre among them. Prenylsynthase commonly contains two aspartate rich motifs, FARM and SARM, responsible for condensing the isoprene units and the deeper cavity depending on the accommodated chain length behind the FARM. Three aspartic acids in the FARM in the prenylsynthase are essential for pyrophosphate recognition. Three aspartic acids D85, D86, and D89 in *Sx*PT1 were conserved with other prenylsynthases, such as 3OYR, and formed the FARM (Figure 1C–D). *Sx*PT1was also composed of a deeper cavity behind the FARM as seen in others (Figure 2A). To estimate the accommodation length of the prenyl chain, the cavity of *Sx*PT1 compared with those of E-PTS (PDB: 3PDE, 29.2% identity with *Sx*PT1), PaFPPS (PDB 3ZOU, 31.9% identity with *Sx*PT1), 3OYR, and *Sinapis alba* GGPP synthase (PDB 2J1P, 31.3% identity with *Sx*PT1) which accommodate DMAPP, GPP, FPP, and GGPP, respectively. Figure 2B-E shows that the cavity of *Sx*PT1 exhibited deeper than PaFPPS, thus *Sx*PT1easily accommodates one molecule of GPP (Figure 2C). Although the superimposing of FPP bound with 3OYR on *Sx*PT1 exhibited the terminus of the prenyl chain clashed with the cavity in *Sx*PT1 (Figure 2D), the cavity elongated the other direction of the superimposed chain and enough volume to place the tip of the chain. It is suggested that *Sx*PT1 may accommodate one molecule of FPP with adapting the chain trace to the shape of the cavity. GGPP seemed to well fit into the cavity of *Sx*PT1 (Figure 2E), contrary to the opposite terminus of the substrate, pyrophosphate, was far from the pyrophosphate recognized residues, D85, D86 and D89 with a distance >10 Å. These comparisons suggested that *Sx*PT1 may recognize and bind to FPP as the longest chain of prenyl donor substrate.

**Figure 2.**
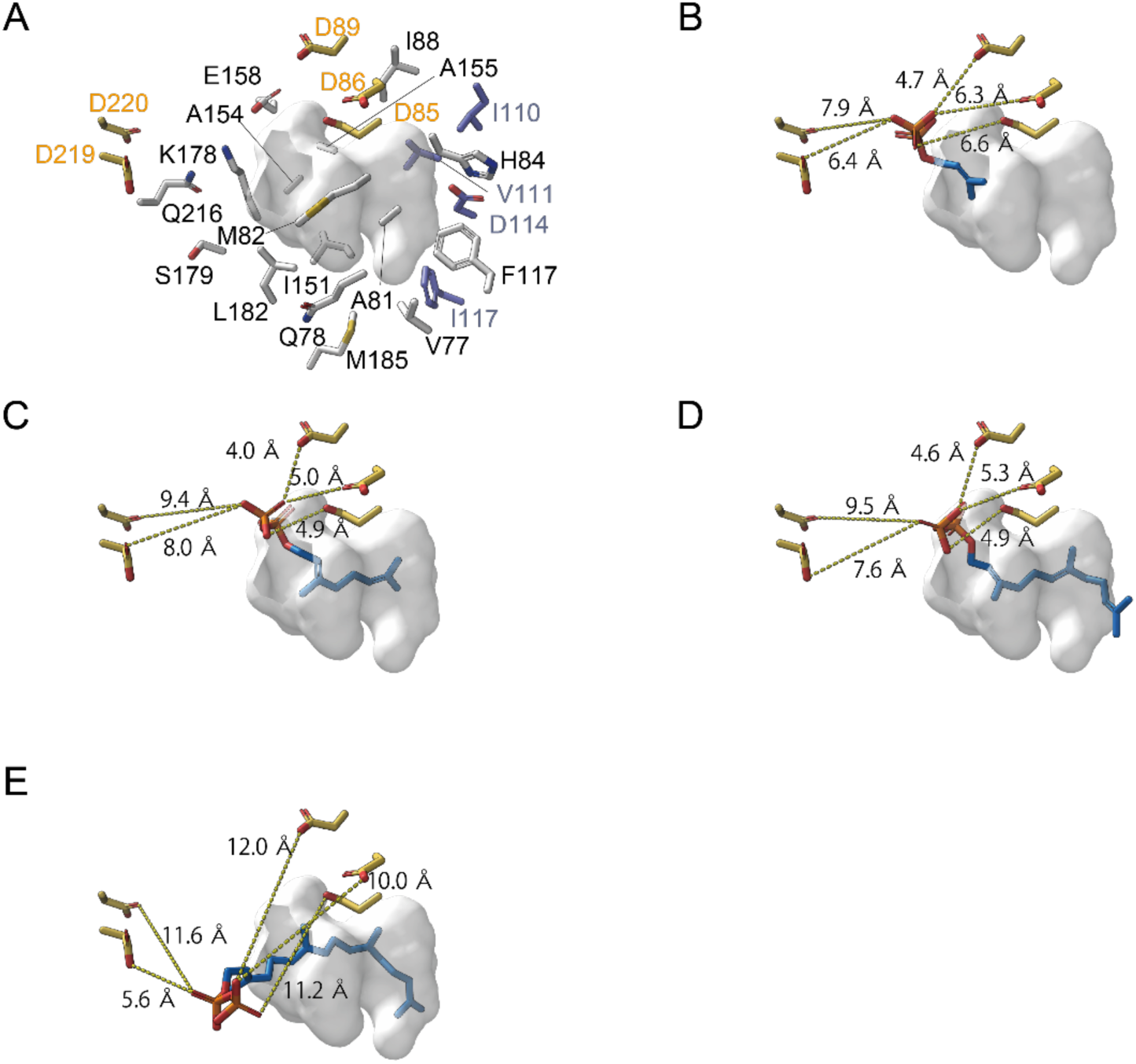
Predicted substrate-accommodation pocket of *Sx*PT1. (A) The predicted substrate-accommodation pocket of *Sx*PT1 was shown as a surface model drawn with a gray color. The sidechains formed by the pocket were shown as sticks. (B) DMAPP bound with E-PTS (PDB 3PDE), (C) GPP bound with PaFPPS (PDB 3ZOU), (D) FPP bound with 3OYR (PDB 3OYR), and (E) GGPP bound with *S. alba* GGPP synthase (PDB 2J1P) were superimposed with *Sx*PT1 and these compounds were shown in the predicted pocket of *Sx*PT1, respectively. The distances between pyrophosphate and Oδ atoms of aspartic acids in ARMs are indicated as yellow dotted lines.

### Complex structure of *Sx*PT1bound with substrate

Based on the structure comparison of *Sx*PT1 with other prenylsynthases, the crystal of apo *Sx*PT1was soaked into the solution of FPP analog (FSPP). The complex structure of *Sx*PT1-FSPP was solved by molecular replacement using apo *Sx*PT1 and consists of two homodimers completely the same as apo *Sx*PT1 with an rmsd value of 0.4 Å (Figure 3A, B). FSPP fit well into the cavity of *Sx*PT1, and the C1 atom of FSPP in *Sx*PT1 was located 3 Å closer to the residue D219 in the pseudo-SARM than that in 3OYR, suggesting that FPP is the longest chain of prenyl donor substrate for *Sx*PT1.

**Figure 3.**
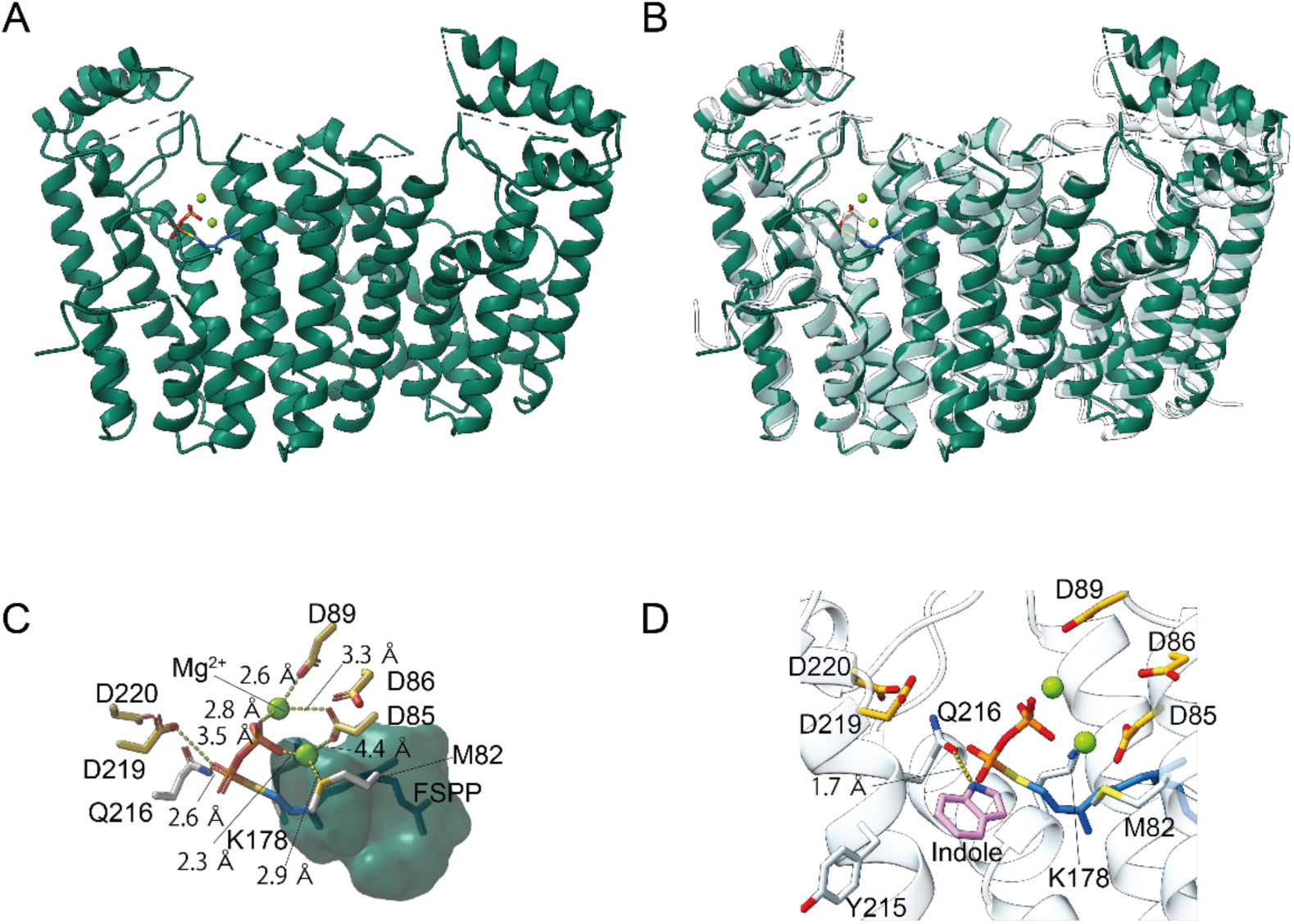
Complex structure of *Sx*PT1 bound with FSPP. (A) The overall structure of *Sx*PT1-FSPP is illustrated as a ribbon diagram. (B) Structural comparison of bound FSPP (green) and apo forms (white) was shown. (C) Close-up view of the substrate-accommodation pocket was shown. The sidechains of aspartic acids and FSPP and Mg^2+^ ions are indicated as sticks and green spheres, respectively The accommodated pocket is indicated as a surface model drawn in green. The D219 in SARM, and D85 and D89 in the FARM interacted directly with FSPP via hydrogen bond and indirectly with FSPP via the coordination of Mg^2+^ ions, respectively. (D) The docking model simulated by MOE was shown. The simulated position of the indole molecule was superimposed onto the complex structure of *Sx*PT1-FSPP. The distances between N1 atoms of indole and Oε atom of Q216 atom is indicated as yellow dotted lines.

Compared to the structures of prenylsynthases previously determined, two Mg^2+^ ions might coordinate with D85 and D89 in the FARM and pyrophosphate (Figure 3C). Additionally, M82 in *Sx*PT1, which is a nonconserved residue, interacted with pyrophosphate via one Mg^2+^ ion. Furthermore, α-phosphate in FSPP was placed closely to the Nε atom of Q216 with 2.6 Å distance to form a hydrogen bond, instead, the interactions between K178 and D86 in FARM and α-phosphate were lost. Although another Mg^2+^ ion coordinated with D222, D223, and D226 in 3OYR and pyrophosphate in DMAPP, no metal ion was found around the residues of D219, D220, A223 in *Sx*PT1 except for a hydrogen bond between Oδ atom of D219 and α-phosphate (Figure 3C).

Structural comparison of the *Sx*PT1 bound with and without FSPP, the sidechain of Y215, which placed closely to the pseudo-SARM, was flipped at 135.5°, creating a space around the pyrophosphate and thus being sufficient to accommodate a prenyl acceptor substrate. According to the sequence similarity of substrate binding site between *Sx*PT1 and other prenyltransferases assigned previously, the interacted and characteristic residues with FSPP in *Sx*PT1, such as Y215, Q216, and pseudo-SARM (D219, D220 and A223), are shared with another prenyltransferase XiaM, predicting that *Sx*PT1 may accommodate an indole substrate as a prenyl acceptor substrate. We tried to soak *Sx*PT1 crystal into the buffer containing with FSPP and indole to obtain the structure of the ternary complex. However, unfortunately, only the binary complex structure of *Sx*PT1-FSPP was observed. Therefore, the docking simulations of indole to the apo *Sx*PT1 were calculated and the simulation predicted the interaction between N1 atom of indole and Oε atom of *Sx*PT1 Q216 via a hydrogen bond (Figure 3D).

### *In vitro* Enzyme reaction of *Sx*PT1

The product of *Sx*PT1 was investigated by the enzyme reaction mixtures to HPLC. The HPLC analysis revealed that the product was present at 38 min in the reaction mixture, but no product was found using boiled *Sx*PT1 (Figure 4A). The enzyme reaction product was determined by LC-MS and exhibited an *m*/*z* value of 322.25299 with an MS tolerance of 1.9 ppm against the theoretical *m*/*z* value of 3-farnesylindole (Figure 4B). To clarify further catalytic role of *Sx*PT1, the product was analyzed using HPLC in the presence of 1-methylindole as a prenyl acceptor substrate (Figure 4B). However, no product in the presence of 1-methylindole was observed by HPLC, suggesting that hydrogen at position N1 atom in the indole is crucial for catalysis by *Sx*PT1.

**Figure 4.**
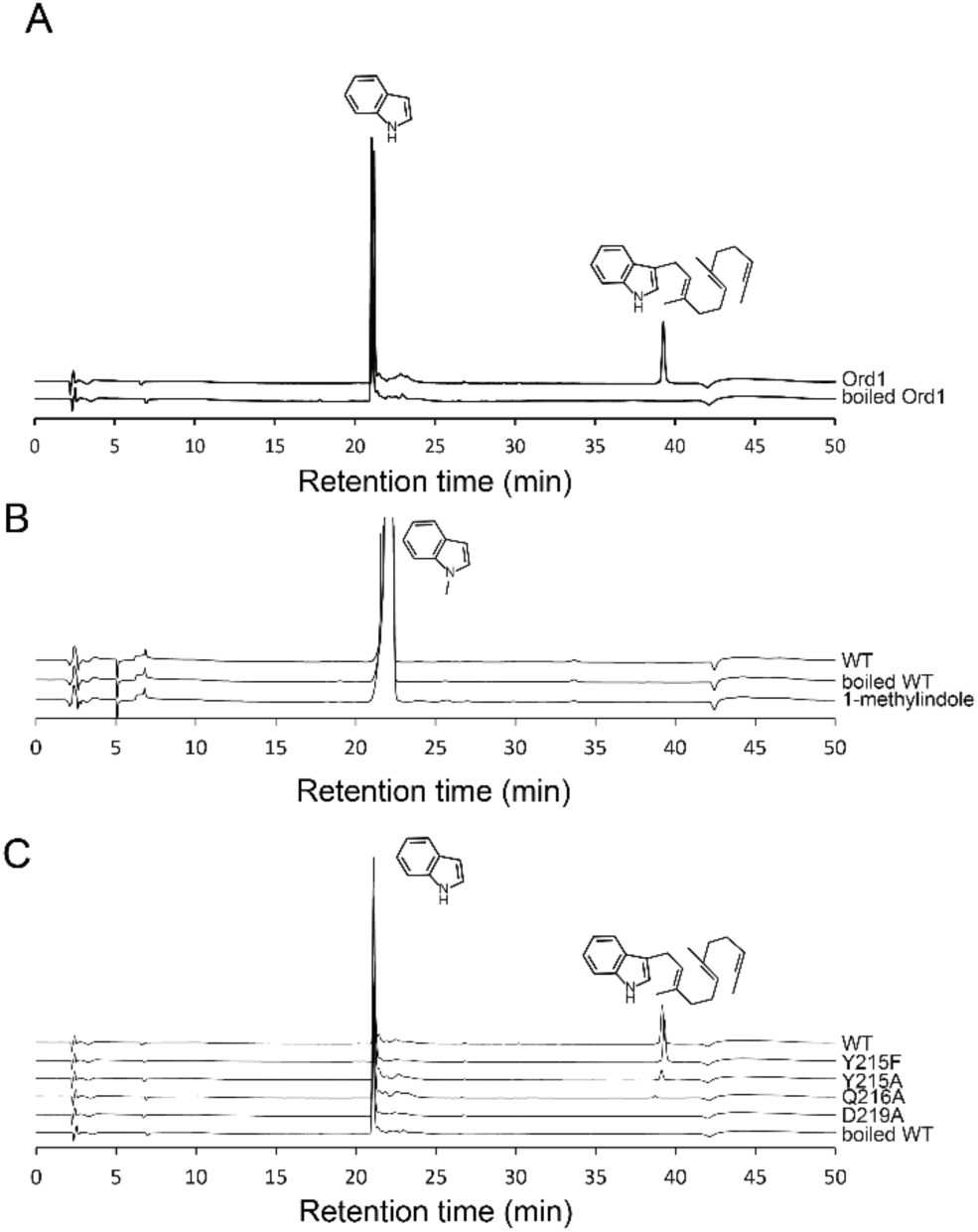
Enzyme reaction product from *Sx*PT1 and its mutant enzymes. Enzyme reaction products from (A) indole and (B) 1-methylindole as the acceptor substrate by *Sx*PT1 and boiled *Sx*PT1 are shown. (C) Enzyme reaction products from indole by *Sx*PT1, Y215F, Y215A, Q216A, and D219A mutant enzymes are shown.

### Structure-based mutagenesis of *Sx*PT1

On the basis of the above observations and simulations, we constructed an *Sx*PT1 site-directed mutant, Q216A. Q216A mutant in the presence of FPP and indole gave the product, however, the yield of the product was significantly decreased with 3% production compared with the wild type, resulting in a nearly complete loss of activity (Figure 4C, Supplementary Table S3). Gln was not deprotonated spontaneously under the enzymatic condition at pH 8.0 consequently, it was not available for abstracting a proton from the N1 atom in the indole directly. The initiation of the catalytic reaction requires a certain active residue to nucleophilic attack to Q216. A detailed analysis of the amino acids around Q216 showed that Oδ atom of D219 was located at one helical pitch downstream from Q216, and was within a distance capable of forming a hydrogen bond with the Nε atom of Q216, then we constructed *Sx*PT1 D219A mutant to clarify whether D219 involved in the catalytic reaction. *In vitro* incubation of D219A mutant mixed with FPP and indole exhibited no product peak appeared (Figure 4C). Since flipping the sidechain of Y215 was observed in the FSPP complex, it anticipated to be necessary for recognizing and accommodating an indole near the reaction site, *Sx*PT1 Y215A and Y215F mutants were also constructed and evaluated their activities. The relative activity of *Sx*PT1 Y215F mutant was 105% compared to that of the wild type.

In contrast to the activity of the Y215F mutant, the relative activity of *Sx*PT1 Y215A compared to the wild type was reduced by over 80% (Figure 4C), suggesting that the aromatic ring at position 215 appropriate position to form a π-π stacking interaction of Tyr/Phe with indole and stabilizes a prenyl acceptor substrate at the proper position.

### Structure of *Sx*PT1 Q216A with substrate and *Sx*PT1 D219A

To confirm the above mutant activities were not caused due to the collapse of the protein structure derived from the substitution of amino acids, the crystal structures of *Sx*PT1 Q216A bound with FSPP and D219A mutants were resolved at resolution 2.2 and 2.6 Å, respectively (Figure 5A–B). Q216A mutant accommodated with FSPP in the cavity as seen in the wild type and was well superimposed with the wild type bound FSPP with an RMSD value of 0.52 Å. Only difference at substitution of the bulkier residue Gln with smaller residue Ala was found (Figure 5C). The coordination manner of Mg^2+^ ions was nearly shared with that in the wild type, i.e., the Mg^2+^ ion was observed only one molecular and coordinated with D85, D89 and the pyrophosphate of FPP (Figure 5C). These results suggested that the structure of Q216A mutant retains its interaction with FSPP independently of the structural change yet failed to produce a product with the indole. It is indicated that the Oε atom of Q216 abstracts the hydrogen from the N1 atom of indole, thereby catalyzing the prenylation. Furthermore, the superimposed RMSD value between *Sx*PT1 D219A mutant and *Sx*PT1 was 0.41 Å (Figure 5B), demonstrating no significant difference was found (Figure 5B and D). Thus, we propose the enzyme reaction mechanism of an all-α-helical prenyltransferase in which the Oδ atom of D219 triggers a nucleophilic attack to initiate the catalytic reaction, leading to the sequential transfer of the prenyl chain to indole (Figure 5E).

**Figure 5.**
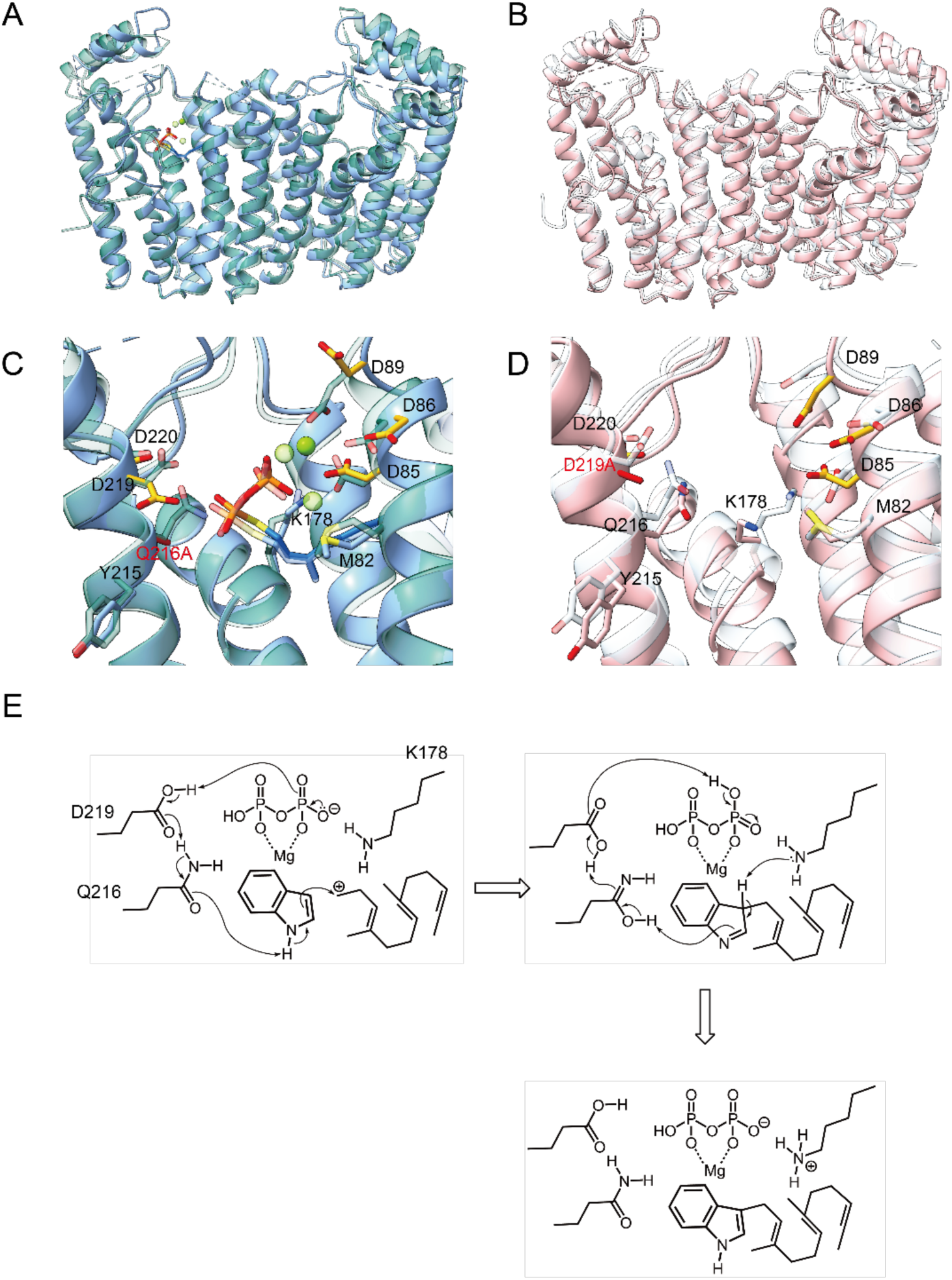
Crystal structures of *Sx*PT1 Q216A and D219A mutants. The overall structures of (A) *Sx*PT1 Q216A-FSPP (blue) superimposed onto *Sx*PT1-FSPP (green) and (B) *Sx*PT1 D219A (pink) superimposed onto *Sx*PT1 (white) are illustrated as ribbon diagrams, respectively. The structural comparison of (C) the *Sx*PT1-FSPP and *Sx*PT1 Q216A-FSPP, and (D) *Sx*PT1 apo and *Sx*PT1 D219A are shown. (E) Predicted catalytic mechanism of *Sx*PT1 was indicated.

## Conclusion

Terpenoids represent the largest and most widely distributed family of natural products^3^. This enormous diversity derived from diverse scaffolds is generated by the prenylsynthase family. In this study, we determined the structure of prenyltransferase *Sx*IPT1 belonging to the all-α-helical prenylsynthase family for the first time. X-ray crystal structures of *Sx*IPT1 and its mutants and their activities revealed that Q216 plays an significant role in interacting N1 atom of indole and Y215 recognizes and places the substrate at the spatial position for the catalytic reaction. Furthermore, these results proposed that Oδ atom of D219 triggers a nucleophilic attack to initiate the catalytic reaction, leading to the Oε atom of Q216 abstracting the hydrogen from the N1 atom of indole and sequential transfer of the prenyl chain to indole. This result may further help the understanding of the catalytic mechanism within all-α-helical prenyltransferase, as well as developing the artificial enzymes based on the prenyltransferase to biosynthesize a wider variety of useful compounds using enzymatic engineering.

## Acknowledgements

This work was supported by JSPS KAKENHI (grant numbers 17KK0141 and 21K05036 to TM). The X-ray diffraction experiments were performed at the Photon Factory (proposals 17G595, 2018G048, 2019G535, 2021G507, and 2023G511). The authors thank FORTE Science Communications (https://www.forte-science.co.jp/) for English language editing.

## Supplementary Material

The Supporting Information is available free of charge on the Publications website.

## Author Contributions

T.I., Y.T., and T.M. designed the study; T.O and S.U. prepared the recombinant proteins, performed structure determination, and evaluated the enzymatic activities; T.O., T.I., Y.K., and T.M. analyzed the data; T.O., T.I., Y.K., T.M. wrote the paper, and T.M. supervised the project.

## References

(1) Sorokina, M.; Steinbeck, C. Review on Natural Products Databases: Where to Find Data in 2020. Journal of Cheminformatics. BioMed Central Ltd. April 3, 2020. 10.1186/s13321-020-00424-9.

(2) Gallo, K.; Kemmler, E.; Goede, A.; Becker, F.; Dunkel, M.; Preissner, R.; Banerjee, P. SuperNatural 3.0 - a Database of Natural Products and Natural Product-Based Derivatives. Nucleic Acids Res 2023, 51 (1 D), D654–D659. 10.1093/nar/gkac1008.

(3) Maczka, W.; Winska, K.; Grabarczyk, M. One Hundred Faces of Geraniol. Molecules. MDPI AG July 1, 2020. 10.3390/molecules25143303.

(4) Xu, B.; Li, Z.; Alsup, T. A.; Ehrenberger, M. A.; Rudolf, J. D. Bacterial Diterpene Synthases Prenylate Small Molecules. ACS Catal 2021, 11 (10), 5906–5915. 10.1021/acscatal.1c01113.

(5) Peralta-Yahya, P. P.; Ouellet, M.; Chan, R.; Mukhopadhyay, A.; Keasling, J. D.; Lee, T. S. Identification and Microbial Production of a Terpene-Based Advanced Biofuel. Nat Commun 2011, 2 (1). 10.1038/ncomms1494.

(6) Klayman, D. L. *Qinghaosu* (Artemisinin): An Antimalarial Drug from China. Science (1979) 1985, 228 (4703), 1049–1055. 10.1126/science.3887571.

(7) Prenyl Transfer Reaction 307. https://pubs.acs.org/sharingguidelines.

(8) Chang, H. Y.; Cheng, T. H.; Wang, A. H. J. Structure, Catalysis, and Inhibition Mechanism of Prenyltransferase. IUBMB Life. Blackwell Publishing Ltd January 1, 2021, pp 40–63. 10.1002/iub.2418.

(9) Liang, P. H.; Ko, T. P.; Wang, A. H. J. Structure, Mechanism and Function of Prenyltransferases. European Journal of Biochemistry. 2002, pp 3339–3354. 10.1046/j.1432-1033.2002.03014.x.

(10) Stepanyuk, A.; Kirschning, A. Synthetic Terpenoids in the World of Fragrances: Iso E Super® Is the Showcase. Beilstein Journal of Organic Chemistry. Beilstein-Institut Zur Forderung der Chemischen Wissenschaften October 31, 2019, pp 2590–2602. 10.3762/bjoc.15.252.

(11) Christianson, D. W. Structural and Chemical Biology of Terpenoid Cyclases. Chemical Reviews. American Chemical Society September 13, 2017, pp 11570–11648. 10.1021/acs.chemrev.7b00287.

(12) Christianson, D. W. Structural Biology and Chemistry of the Terpenoid Cyclases. Chem Rev 2006, 106 (8), 3412–3442. 10.1021/cr050286w.

(13) Chan, Y. Te; Ko, T. P.; Yao, S. H.; Chen, Y. W.; Lee, C. C.; Wang, A. H. J. Crystal Structure and Potential Head-to-Middle Condensation Function of a Z,Z-Farnesyl Diphosphate Synthase. ACS Omega 2017, 2 (3), 930–936. 10.1021/acsomega.6b00562.

(14) Christianson, D. W. Structural Biology and Chemistry of the Terpenoid Cyclases. Chem Rev 2006, 106 (8), 3412–3442. 10.1021/cr050286w.

(15) Tello, M.; Kuzuyama, T.; Heide, L.; Noel, J. P.; Richard, S. B. The ABBA Family of Aromatic Prenyltransferases: Broadening Natural Product Diversity. Cellular and Molecular Life Sciences. May 2008, pp 1459–1463. 10.1007/s00018-008-7579-3.

(16) Mai, P.; Zocher, G.; Stehle, T.; Li, S. M. Structure-Based Protein Engineering Enables Prenyl Donor Switching of a Fungal Aromatic Prenyltransferase. Org Biomol Chem 2018, 16 (40), 7461– 7469. 10.1039/C8OB02037J.

(17) Qian, S.; Clomburg, J. M.; Gonzalez, R. Engineering Escherichia Coli as a Platform for the in Vivo Synthesis of Prenylated Aromatics. Biotechnol Bioeng 2019, 116 (5), 1116–1127. 10.1002/bit.26932.

(18) Huang, H.; Levin, E. J.; Liu, S.; Bai, Y.; Lockless, S. W.; Zhou, M. Structure of a Membrane-Embedded Prenyltransferase Homologous to UBIAD1. PLoS Biol 2014, 12 (7), 1–11. 10.1371/journal.pbio.1001911.

(19) Huang, H.; Scherman, M. S.; D’Haeze, W.; Vereecke, D.; Holsters, M.; Crick, D. C.; McNeil, M. R. Identification and Active Expression of the Mycobacterium Tuberculosis Gene Encoding 5-Phospho-α-D-Ribose-1-Diphosphate: Decaprenyl-Phosphate 5-Phosphoribosyltransferase, the First Enzyme Committed to Decaprenylphosphoryl-D-Arabinose Synthesis. Journal of Biological Chemistry 2005, 280 (26), 24539–24543. 10.1074/jbc.M504068200.

(20) Huang, H.; Levin, E. J.; Liu, S.; Bai, Y.; Lockless, S. W.; Zhou, M. Structure of a Membrane-Embedded Prenyltransferase Homologous to UBIAD1. PLoS Biol 2014, 12 (7), 1–11. 10.1371/journal.pbio.1001911.

(21) Winkelblech, J.; Fan, A.; Li, S. M. Prenyltransferases as Key Enzymes in Primary and Secondary Metabolism. Applied Microbiology and Biotechnology. Springer Verlag September 22, 2015, pp 7379–7397. 10.1007/s00253-015-6811-y.

(22) Wang, J.; Chu, S.; Zhu, Y.; Cheng, H.; Yu, D. Positive Selection Drives Neofunctionalization of the UbiA Prenyltransferase Gene Family. Plant Mol Biol 2015, 87 (4–5), 383–394. 10.1007/s11103-015-0285-2.

(23) Ding, L.; Münch, J.; Goerls, H.; Maier, A.; Fiebig, H. H.; Lin, W. H.; Hertweck, C. Xiamycin, a Pentacyclic Indolosesquiterpene with Selective Anti-HIV Activity from a Bacterial Mangrove Endophyte. Bioorg Med Chem Lett 2010, 20 (22), 6685–6687. 10.1016/j.bmcl.2010.09.010.

(24) Pfaffenbach, M.; Bakanas, I.; O’Connor, N. R.; Herrick, J. L.; Sarpong, R. Total Syntheses of Xiamycins A, C, F, H and Oridamycin A and Preliminary Evaluation of Their Anti-Fungal Properties. Angewandte Chemie 2019, 131 (43), 15448–15452. 10.1002/ange.201908399.

(25) Miyata, A.; Ito, S.; Fujinami, D. Structure Prediction and Genome Mining-Aided Discovery of the Bacterial C-Terminal Tryptophan Prenyltransferase PalQ. Advanced Science 2024, 11 (6). 10.1002/advs.202307372.

(26) McIntosh, J. A.; Donia, M. S.; Nair, S. K.; Schmidt, E. W. Enzymatic Basis of Ribosomal Peptide Prenylation in Cyanobacteria. J Am Chem Soc 2011, 133 (34), 13698–13705. 10.1021/ja205458h.

(27) Zhang, Y.; Goto, Y.; Suga, H. Discovery, Biochemical Characterization, and Bioengineering of Cyanobactin Prenyltransferases. Trends in Biochemical Sciences. Elsevier Ltd April 1, 2023, pp 360–374. 10.1016/j.tibs.2022.11.002.

(28) Luk, L. Y. P.; Tanner, M. E. Mechanism of Dimethylallyltryptophan Synthase: Evidence for a Dimethylallyl Cation Intermediate in an Aromatic Prenyltransferase Reaction. J Am Chem Soc 2009, 131 (39), 13932–13933. 10.1021/ja906485u.

(29) Hao, Y.; Pierce, E.; Roe, D.; Morita, M.; McIntosh, J. A.; Agarwal, V.; Cheatham, T. E.; Schmidt, E. W.; Nair, S. K. Molecular Basis for the Broad Substrate Selectivity of a Peptide Prenyltransferase. Proc Natl Acad Sci U S A 2016, 113 (49), 14037–14042. 10.1073/pnas.1609869113.

(30) Okada, M.; Ishihara, A.; Yamasaki, R.; Tsuji, F.; Hayashi, S.; Usami, S.; Sakagami, Y. A Region Corresponding to Second Aspartate-Rich Motif in Tryptophan Isoprenylating Enzyme, ComQ, Serves as a Substrate-Binding Site. Biosci Biotechnol Biochem 2014, 78 (4), 550–555. 10.1080/09168451.2014.891932.

(31) Hirooka, K.; Shioda, S.; Okada, M. Identification of Critical Residues for the Catalytic Activity of ComQ, a Bacillus Prenylation Enzyme for Quorum Sensing, by Using a Simple Bioassay System. Biosci Biotechnol Biochem 2020, 84 (2), 347–357. 10.1080/09168451.2019.1685371.

(32) Ito, H.; Matsui, T.; Konno, R.; Itakura, M.; Kodera, Y. LC–MS Peak Assignment Based on Unanimous Selection by Six Machine Learning Algorithms. Sci Rep 2021, 11 (1). 10.1038/s41598-021-02899-4.

(33) Rice, L. M.; Earnest, T. N.; Brunger, A. T.; Howard, C. Biological Crystallography Single-Wavelength Anomalous Diffraction Phasing Revisited. Acta Cryst 2000.

(34) Kabsch, W. XDS. Acta Crystallogr D Biol Crystallogr 2010, 66 (2), 125–132. 10.1107/S0907444909047337.

(35) Terwilliger, T. C.; Adams, P. D.; Read, R. J.; McCoy, A. J.; Moriarty, N. W.; Grosse-Kunstleve, R. W.; Afonine, P. V.; Zwart, P. H.; Hung, L. W. Decision-Making in Structure Solution Using Bayesian Estimates of Map Quality: The PHENIX AutoSol Wizard. Acta Crystallogr D Biol Crystallogr 2009, 65 (6), 582–601. 10.1107/S0907444909012098.

(36) Terwilliger, T. C.; Grosse-Kunstleve, R. W.; Afonine, P. V.; Moriarty, N. W.; Zwart, P. H.; Hung, L. W.; Read, R. J.; Adams, P. D. Iterative Model Building, Structure Refinement and Density Modification with the PHENIX AutoBuild Wizard. In Acta Crystallographica Section D: Biological Crystallography; 2007; Vol. 64, pp 61–69. 10.1107/S090744490705024X.

(37) Vagin, A.; Teplyakov, A. MOLREP: An Automated Program for Molecular Replacement; 1997; Vol. 30.

(38) Emsley, P.; Lohkamp, B.; Scott, W. G.; Cowtan, K. Features and Development of Coot. Acta Crystallogr D Biol Crystallogr 2010, 66 (4), 486–501. 10.1107/S0907444910007493.

(39) Afonine, P. V.; Grosse-Kunstleve, R. W.; Echols, N.; Headd, J. J.; Moriarty, N. W.; Mustyakimov, M.; Terwilliger, T. C.; Urzhumtsev, A.; Zwart, P. H.; Adams, P. D. Towards Automated Crystallographic Structure Refinement with *Phenix.Refine*. Acta Crystallogr D Biol Crystallogr 2012, 68 (4), 352–367. 10.1107/S0907444912001308.

(40) Williams, C. J.; Headd, J. J.; Moriarty, N. W.; Prisant, M. G.; Videau, L. L.; Deis, L. N.; Verma, V.; Keedy, D. A.; Hintze, B. J.; Chen, V. B.; Jain, S.; Lewis, S. M.; Arendall, W. B.; Snoeyink, J.; Adams, P. D.; Lovell, S. C.; Richardson, J. S.; Richardson, D. C. MolProbity: More and Better Reference Data for Improved All-Atom Structure Validation. Protein Science 2018, 27 (1), 293–315. 10.1002/pro.3330.

(41) Schrödinger, L.; DeLano, W. PyMOL. http://www.pymol.org/pymol (accessed 2023-01-18).

(42) Pettersen, E. F.; Goddard, T. D.; Huang, C. C.; Couch, G. S.; Greenblatt, D. M.; Meng, E. C.; Ferrin, T. E. UCSF Chimera - A Visualization System for Exploratory Research and Analysis. J Comput Chem 2004, 25 (13), 1605–1612. 10.1002/jcc.20084.

(43) Holm, L.; Laiho, A.; Törönen, P.; Salgado, M. DALI Shines a Light on Remote Homologs: One Hundred Discoveries. Protein Science 2023, 32 (1). 10.1002/pro.4519.

(44) Madeira, F.; Madhusoodanan, N.; Lee, J.; Eusebi, A.; Niewielska, A.; Tivey, A. R. N.; Lopez, R.; Butcher, S. The EMBL-EBI Job Dispatcher Sequence Analysis Tools Framework in 2024. Nucleic Acids Res 2024, 52 (W1), W521–W525. 10.1093/nar/gkae241.

(45) Robert, X.; Gouet, P. Deciphering Key Features in Protein Structures with the New ENDscript Server. Nucleic Acids Res 2014, 42 (W1). 10.1093/nar/gku316.

