## Supplementary materials for "Structure-activity relationship of an all-α-helical prenyltransferase reveals a mechanism for indole prenylation"

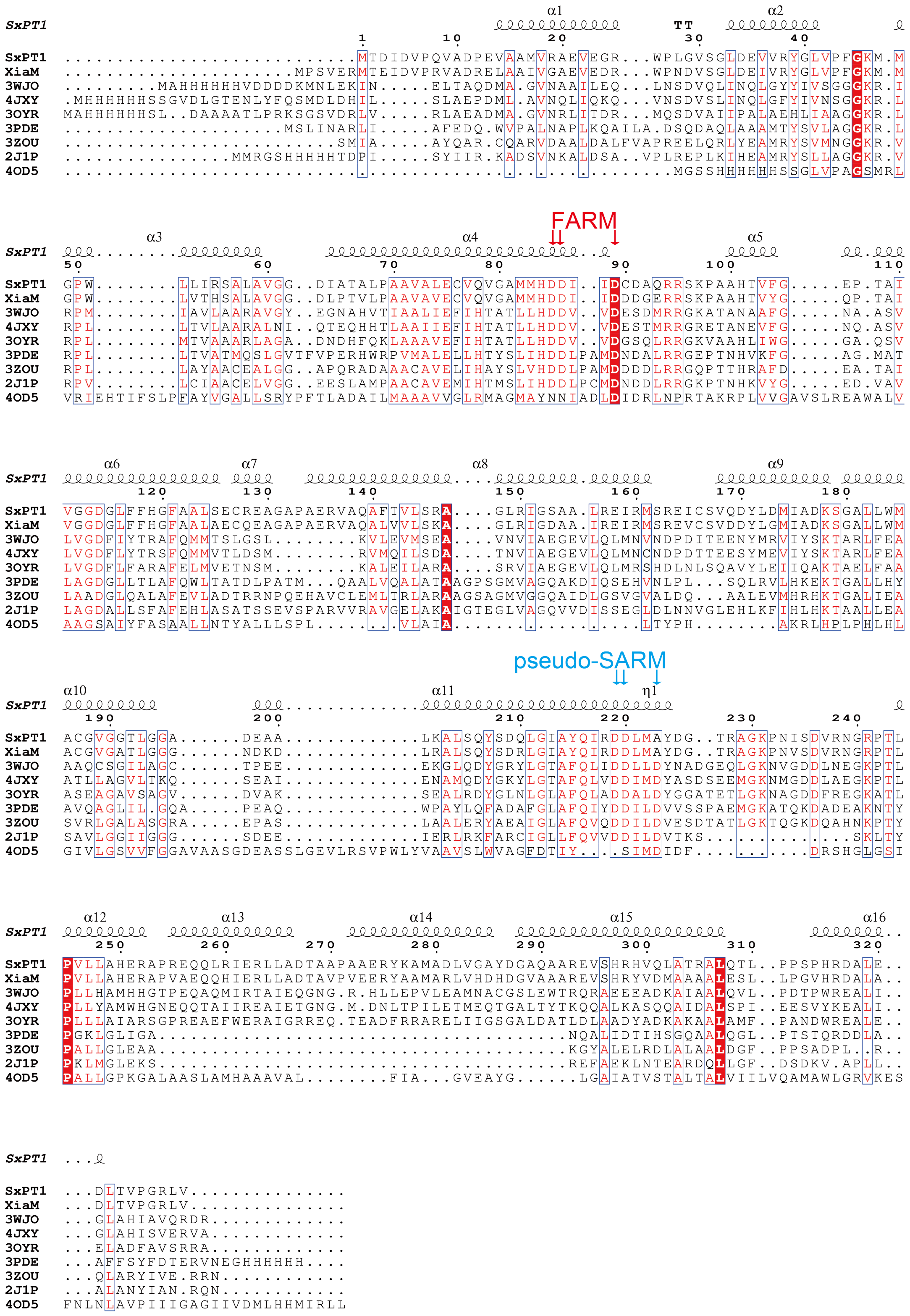

α1
α2
SxPT1
TT
1 10 20 30 40
SxPT1 ...........................MTDIDVPQVADPEVAAMVRAEVEGR...WPLGVSGLDEVVRYGLVPFGKM.M
XiaM .....................MPSVERMTEIDVPRVADRELAAIVGAEVEDR...WPNDVSGLDEIVRYGLVPFGKM.M
3WJO .........MAHHHHHHVDDDDKMNLEKIN.....ELTAQDMA.GVNAAILEQ...LNSDVQLINQLGYYIVSGGGKR.I
4JXY .MHHHHHHSSGVDLGTENLYFQSMDLDHIL.....SLAEPDML.AVNQLIQKQ...VNSDVSLINQLGFYIVNSGGKR.L
3OYR MAHHHHHHSL..DAAATLPRKSGSVDRLV.....RLAEADMA.GVNRLITDR...MQSDVAIIPALAEHLIAAGGKR.L
3PDE ...................MSLINARLI.....AFEDQ.WVPALNAPLKQAILA.DSQDAQLAAAMTYSVLAGGKR.L
3ZOU .........................SMIA.....AYQAR.CQARVDAALDALFVAPREELQRLYEAMRYSVMNGGKR.V
2J1P ...............MMRGSHHHHHHTDPI.....SYIIR.KADSVNKALDSA..VPLREPLKIHEAMRYSLLAGGKR.V
4OD5 ..................................................MGSSHHHHHSSGLVPAGSMRL
FARM
α3
α4
α5
50 60 70 80 90 100 110
SxPT1 GPW........LLIRSALAVGG..DIATALPAAVALECVQVGAMMHDDI..IDCDAQRRSKPAAHTVFG.....EP.TAI
XiaM GPW........LVTHSALAVGG..DLPTVLPAAVAVECVQVGAMMHDDI..IDDDGERRSKPAAHTVYG.....QP.TAI
3WJO RPM........IAVLAARAVGY..EGNAHVTIAALIEFIHTATLLHDDV..VDESDMRRGKATANAAFG.....NA.ASV
4JXY RPL........LTVLAARALNI..QTEQHHTLAAIIEFIHTATLLHDDV..VDESTMRRGRETANEVFG.....NQ.ASV
3OYR RPL........MTVAAARLAGA..DNDHFQKLAAAVEFIHTATLLHDDV..VDGSQLRRGKVAAHLIWG.....GA.QSV
3PDE RPL........LTVATMQSLGVTFVPERHWRPVMALELLHTYSLIHDDLPAMDNDALRRGEPTNHVKFG.....AG.MAT
3ZOU RPL........LAYAACEALGG..APQRADAAACAVELIHAYSLVHDDLPAMDDDDLRRGQPTTHRAFD.....EA.TAI
2J1P RPV........LCIAACELVGG..EESLAMPAACAVEMIHTMSLIHDDLPCMDNDDLRRGKPTNHKVYG.....ED.VAV
4OD5 VRIEHTIFSLPFAYVGALLSRYPFTLADAILMAAAVVGLRMAGMAYNNIADLDIDRLNPRTAKRPLVVGAVSLREAWALV
α6
α7
α8
α9
120 130 140 150 160 170 180
SxPT1 VGGDGLFFHGFAALSECREAGAPAERVAQAFTVLSRA....GLRIGSAA.LREIRMSREICSVQDYLDMIADKSGALLWM
XiaM VGGDGLFFHGFAALAECQEAGAPAERVAQALVVLSKA....GLRIGDAA.IREIRMSREVCSVDDYLGMIADKSGALLWM
3WJO LVGDFIYTRAFQMMTSLGSL........KVLEVMSEA....VNVIAEGEVLQLMNVNDPDITEENYMRVIYSKTARLFEA
4JXY LVGDFLYTRSFQMMVTLDSM........RVMQILSDA....TNVIAEGEVLQLMNCNDPDTTEESYMEVIYSKTARLFEA
3OYR LVGDFLFARAFELMVETNSM........KALEILARA....SRVIAEGEVLQLMRSHDLNLSQAVYLEIIQAKTAELFAA
3PDE LAGDGLLTLAFQWLTATDLPATM...QAALVQALATAAGPSGMVAGQAKDIQSEHVNLPL...SQLRVLHKEKTGALLHY
3ZOU LAADGLQALAFEVLADTRRNPQEHAVCLEMLTRLARAAGSAGMVGGQAIDLGSVGVALDQ...AALEVMHRHKTGALIEA
2J1P LAGDALLSFAFEHLASATSSEVSPARVVRAVGELAKAIGTEGLVAGQVVDISSEGLDLNNVGLEHLKFIHLHKTAALLEA
4OD5 AAGSAIYFASAALLNTYALLLSPL.........VLAIA...........................LTYPH........AKRLHPLPHLHL
pseudo-SARM
α10
α11
η1
190 200 210 220 230 240
SxPT1 ACGVGGTLGGA......DEAA.............LKALSQYSDQLGIAYQIRDDLMAYDG..TRAGKPNISDVRNGRPTL
XiaM ACGVGATLGGG......NDKD.............LRALSQYSDRLGIAYQIRDDLMAYDG..TRAGKPNVSDVRNGRPTL
3WJO AAQCSGILAGC......TPEE.............EKGLQDYGRYLGTAFQLIDDLLDYNADGEQLGKNVGDDLNEGKPTL
4JXY ATLLAGVLTKQ......SEAI.............ENAMQDYGKYLGTAFQLVDDIMDYASDSEEMGKNMGDDLAEGKPTL
3OYR ASEAGAVSAGV......DVAK.............SEALRDYGLNLGLAFQLADDALDYGGATETLGKNAGDDFREGKATL
3PDE AVQAGLIL.GQA.....PEAQ.............WPAYLQFADAFGLAFQIYDDILDVVSSPAEMGKATQKDADEAKNTY
3ZOU SVRLGALASGRA.....EPAS.............LAALERYAEAIGLAFQVQDDILDVESDTATLGKTQGKDQAHNKPTY
2J1P SAVLGGIIGGG......SDEE.............IERLRKFARCIGLLFQVVDDILDVTKS...............SKLTY
4OD5 GIVLGSVVFGGAVAASGDEASSLGEVLRSVPWLYVAAVSLWVAGFDTIY....SIMDIDF...............DRSHGLGSI
α12
α13
α14
α15
α16
250 260 270 280 290 300 310 320
SxPT1 PVLLAHERAPREQQLRIERLLADTAAPAAERYKAMADLVGAYDGAQAAREVSHRHVQLATRALQTL..PPSPHRDALE..
XiaM PVLLAHERAPVAEQQHIERLLADTAVPVEERYAAMARLVHDHDGVAAAREVSHRYVDMAAAALESL..LPGVHRDALA..
3WJO PLLHAMHHGTPEQAQMIRTAIEQGNG.R.HLLEPVLEAMNACGSLEWTRQRAEEEADKAIAALQVL..PDTPWREALI..
4JXY PLLYAMWHGNEQQTAIIREAIETGNG.M.DNLTPILETMEQTGALTYTKQQALKASQQAIDALSPI..EESVYKEALI..
3OYR PLLLAIARSGPREAEFWERAIGRREQ.TEADFRRARELIIGSGALDATLDLAADYADKAKAALAMF..PANDWREALE..
3PDE PGKLGLIGA...................................NQALIDTIHSGQAALQGL..PTSTQRDDLA..
3ZOU PALLGLEAA...................................KGYALELRDLALAALDGF..PPSADPL..R..
2J1P PKLMGLEKS...................................REFAEKLNTEARDQLLGF..DSDKV.APLL..
4OD5 PALLGPKGALAASLAMHAAAVAL........FIA...GVEAYG......LGAIATVSTALTALVIILVQAMAWLGRVKES
SxPT1 ...
SxPT1 ...DLTVPGRLV...............
XiaM ...DLTVPGRLV...............
3WJO ...GLAHIAVQRDR.............
4JXY ...GLAHISVERVA.............
3OYR ...ELADFAVSRRA.............
3PDE ...AFFSYFDTERVNEGHHHHHH....
3ZOU ...QLARYIVE.RRN............
2J1P ...ALANYIAN.RQN............
4OD5 FNLNLAVPIIIGAGIIVDMLHHMIRLL


17

**Supplementary Figure S1**. Multiple sequence alignment of prenyltransferases and prenylsynthases.

Sequence alignment of two all-α-prenyltransferases (SxPT1 and XiaM), six prenylsynthases (3WJO, 4JXY, 3OYR, 3PDE, 3ZOU), and Ubi-A prenyltransferase using clustal OMEGA (1.2.4)[44], visualized with ESPript 3.0[45] (https://endscript.ibcp.fr). Conserved sequences colored red. Secondary structure elements as identified in the crystal structure of *Sx*PT1 molA. FARM and pseudo-SARM sequences are indicated by red and blue arrow respectively. XiaM, 3WJO, 4JXY, 3OYR, 3PDE, 3ZOU, 2J1P share 79.6, 26.6, 27.6, 30.4, 29.2, 31.9, and 31.3% identities with *Sx*PT1 respectively. 4OD5 was not found significant similarity with *Sx*PT1.

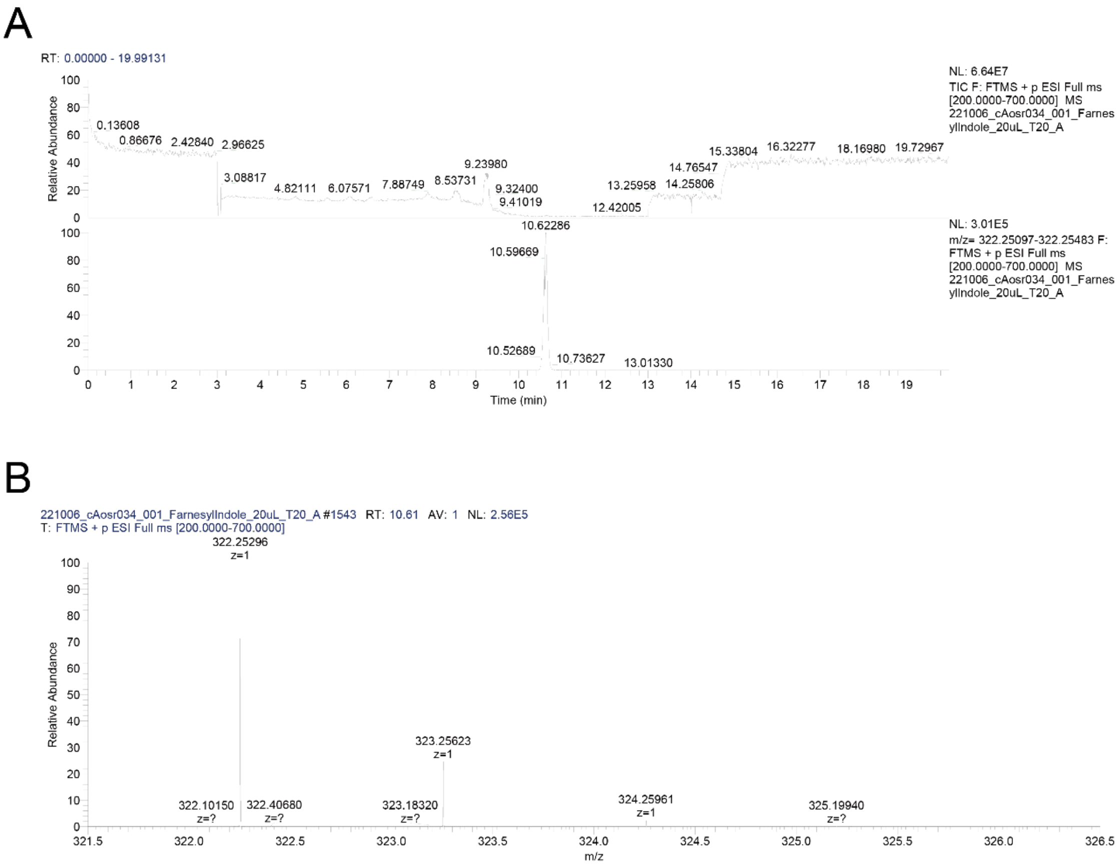


**Supplementary Figure S2**. LC–MS analysis of the products of *Sx*PT1 enzyme assays.

(A) Total ion chromatogram ranging from 200- 700 *m/z* was shown (top panel). Extracted ion chromatograms at m/z value of 322.25299 with a mass tolerance 6 ppm was shown (lower panel). (B) Mass spectrum of the peaks at retention time of 10.61 min was shown.

**Supplementary Table S1**. Primers for *Sx*PT1 mutations.

| Primer name | Sequence (5' to 3') |
|---|---|
| Y215A-F | AGCTGGGCATCGCCGCCCAGATCCGTGACG |
| Y215A-R | CGTCACGGATCTGGGCGGCGATGCCCAGCT |
| Y215F-F | AGCTGGGCATCGCCTTCCAGATCCGTGACG |
| Y215F-R | CGTCACGGATCTGGAAGGCGATGCCCAGCT |
| Q216A-F | TGGGCATCGCCTACGCGATCCGTGACGACC |
| Q216A-R | GGTCGTCACGGATCGCGTAGGCGATGCCCA |
| D219A-F | ACGCCATGAGGTCGGCACGGATCTGGTAG |
| D219A-R | CTACCAGATCCGTGCCGACCTCATGGCGT |

The sequence belonging to the mutate sequence is underlined.

**Supplementary Table S2**. Summary of crystallization conditions, data-collection and refinement statistics[a].

| | *Sx*PT1 SeMet | *Sx*PT1 | *Sx*PT1 - FSPP | *Sx*PT1 Q216A-FSPP | *Sx*PT1 D219A |
|---|---|---|---|---|---|
| **Data collection** | | | | | |
| Beamline | PF BL-5A | PF BL-1A | PF BL-1A | PF BL-1A | PF BL-17A |
| Wavelength (Å) | 0.97922 | 1.04500 | 1.04500 | 1.04500 | 0.98000 |
| Space group | *P*1 | *P*1 | *P*1 | *P*1 | *P*1 |
| a, b, c (Å) | 59.3, 72.7, 84.3 | 59.0, 72.2, 84.2 | 59.2, 72.6, 84.4 | 58.9, 72.2, 84.0 | 58.9, 72.4, 83.8 |
| $\alpha$, β, γ (°) | 94.3, 69.5, 90.0 | 86.0, 69.5, 90.0 | 93.9, 69.5, 90.1 | 86.2, 69.4, 89.9 | 93.5, 110.5, 90.1 |
| Resolution range (Å) | 50.0-1.78 (1.89-1.78) | 50.0-2.10 (2.22-2.10) | 50.0-2.00 (2.12-2.00) | 50.00-1.95 (2.07-1.95) | 50.0-2.80 (2.97-2.80) |
| Completeness (%) | 95.8 (93.7) | 94.6 (95.0) | 97.6 (96.6) | 97.5 (96.4) | 94.8 (94.7) |
| $<I/\sigma(I)>$ | 31.53 (2.85) | 7.71 (1.66) | 12.71 (1.75) | 13.58 (2.05) | 7.53 (1.94) |
| $R_{merge}$[b] (%) | 3.0 (66.8) | 7.6 (61.6) | 5.2 (70.6) | 6.2 (64.6) | 10.5 (64.4) |
| $CC_{1/2}$[c] | 1.000 (0.880) | 0.997 (0.667) | 0.999 (0.746) | 0.999 (0.824) | 0.993 (0.433) |
| Multiplicity | 7.2 (7.4) | 3.4 (3.0) | 3.6 (3.7) | 3.6 (3.7) | 1.8 (1.9) |
| Total reflections | 1,739,580 (281,067) | 267,449 (44,693) | 312,316 (51,426) | 333,550 (54,765) | 110,549 (18,080) |
| No. of unique reflections | 242,171 (38,269) | 144,033 (23,415) | 86,258 (13,824) | 91,888 (14,660) | 60,573 (9,723) |
| **Refinement collection** | | | | | |
| Resolution range (Å) | 44.75-1.78 | 44.40-2.10 | 55.62-2.15 | 44.26-2.20 | 44.35-2.80 |
| $R_{work}$[d] (%) | 26.67 | 23.08 | 21.08 | 22.39 | 23.35 |
| $R_{free}$[e] (%) | 28.44 | 24.86 | 25.82 | 24.38 | 26.90 |
| RMS (bonds) (Å) | 0.008 | 0.003 | 0.008 | 0.003 | 0.03 |
| RMS (angles) (°) | 1.057 | 0.668 | 1.049 | 0.591 | 0.669 |
| Twin fraction | 0.500 (*h, -k, h-l*) | 0.500 (*h, -k, h-l*) | 0.500 (*h, -k, h-l*) | 0.500 (*h, -k, h-l*) | 0.500 (*h, -k, h-l*) |
| Clash score | 13.53 | 8.89 | 10.06 | 8.49 | 12.92 |
| **Ramachandran plot** | | | | | |
| Ramachandran favored (%) | | 98.66 | 98.95 | 98.26 | 98.75 |
| Ramachandran allowed (%) | | 1.35 | 0.86 | 1.74 | 1.25 |
| Ramachandran outliers (%) | | 0.09 | 0.19 | 0 | 0 |

[a] Values in parentheses are for the highest resolution shell.

[b] $R_{merge} = \Sigma_h \Sigma_j |<I>_h - I_{h,j}|/\Sigma_h \Sigma_j I_{h,j}$, where $<I>_h$ is the mean intensity of symmetry-equivalent reflections.

[c] $CC_{1/2}$ is the Pearson correlation coefficient for a random half of the data, the two numbers represent the lowest and highest resolution shell, respectively.

**Supplementary Table S3**. Relative activities of the SxPT1 and mutants.

| Mutants | Relative activity (%) |
|---|---|
| WT | 100.0 ± 5.1 |
| Y215F | 104.9 ± 7.5 |
| Y215A | 11.2 ± 1.4 |
| Q216A | 4.2 ± 0.2 |
| D215A | N/D |

Values are means ± standard deviation (n = 3). N/D means not detected

[d] $R_{work} = \Sigma|F_{obs} - F_{cal}|/\Sigma F_{obs}$, where $F_{obs}$ and $F_{cal}$ are observed and calculated structure factor amplitudes, respectively.

[e] $R_{free}$ value was calculated for $R$ factor, using only an unrefined subset of reflection data.
